# Inhibition of noradrenaline-dependent synaptic transmission in the dorsal raphe nucleus by alpha2-adrenergic receptors

**DOI:** 10.1101/2023.11.07.566093

**Authors:** Aleigha Gugel, Erik A. Ingebretsen, Holly S. Hake, Stephanie C. Gantz

## Abstract

In the central nervous system, noradrenaline transmission controls the degree to which we are awake, alert, and attentive. Aberrant noradrenaline transmission is associated with pathological forms of hyper- and hypo-arousal that present in numerous neuropsychiatric disorders often associated with dysfunction in serotonin transmission. *In vivo,* noradrenaline regulates the release of serotonin because noradrenergic input drives the serotonin neurons to fire action potentials via activation of excitatory α1-adrenergic receptors (α1-A_R_). Despite the critical influence of noradrenaline on the activity of dorsal raphe serotonin neurons, the source of noradrenergic afferents has not been resolved and the presynaptic mechanisms that regulate noradrenaline-dependent synaptic transmission have not been described. Using an acute brain slice preparation from male and female mice and electrophysiological recordings from dorsal raphe serotonin neurons, we found that selective optogenetic activation of locus coeruleus terminals in the dorsal raphe was sufficient to produce an α1-A_R_-mediated excitatory postsynaptic current (α1-A_R_-EPSC). Activation of inhibitory α2-adrenergic receptors (α2-A_R_) with UK-14,304 eliminated the α1-A_R_-EPSC via presynaptic inhibition of noradrenaline release, likely via inhibition of voltage-gated calcium channels. In a subset of serotonin neurons, activation of postsynaptic α2-A_R_ produced an outward current through activation of potassium conductance. Further, *in vivo* activation of α2-A_R_ by systemic administration of clonidine reduced the expression of c-fos in the dorsal raphe serotonin neurons, indicating reduced neural activity. Thus, α2-A_R_ are critical regulators of serotonin neuron excitability.

## Introduction

In the central nervous system, noradrenaline transmission controls the degree to which we are awake, alert, and attentive. Action potential firing of the noradrenaline neurons in the locus coeruleus (LC) drives the transition from sleep to wake. Once awake, LC neurons are excited by salient stimuli, including those that are non-stressful as well as physiological and psychosocial stressors (reviewed in [1]). An elevation in the rate of LC neuron firing causes increased noradrenaline release in terminal projection areas, ultimately leading to heightened arousal which promotes specific motor behaviors that aid in survival. However, aberrant noradrenaline transmission is associated with pathological forms of hyper- and hypo-arousal in sleep disturbances, pain syndromes, post-traumatic stress (PTSD), attention deficit hyperactivity (ADHD), obsessive-compulsive, bipolar, schizophrenia, depression, and anxiety disorders. Notably, these disorders are also associated with dysregulation of serotonin transmission. Consequently, altering the levels of noradrenaline and serotonin in the brain is a major pharmacological target to treat neuropsychiatric disorders. Gaining a comprehensive understanding of the mechanisms by which these monoamine systems interact is necessary to develop more efficacious therapies.

The principal source of serotonin released throughout the brain is the population of the serotonin neurons that reside in the dorsal raphe nucleus. *In vivo,* the levels of noradrenaline and serotonin in the dorsal raphe covary positively [2]. This covariance arises because noradrenergic input drives the serotonin neurons to fire action potentials and release serotonin via activation of Gαq protein-coupled α1-adrenergic receptors (α1-A_R_) [3–9]. In brain slices containing the dorsal raphe, tonic noradrenergic input is severed but action potential firing in the serotonin neurons can be restored with either exogenous α1-A_R_ agonists [10] or by electrically stimulating the vesicular release of noradrenaline which evokes an α1-A_R_-dependent excitatory postsynaptic current (α1-A_R_-EPSC) [11,12]. Despite the critical influence of noradrenaline on the activity of dorsal raphe serotonin neurons, the source of noradrenergic afferents has not been resolved. Furthermore, the presynaptic mechanisms that regulate noradrenaline-dependent synaptic transmission have not been described.

Using whole-cell patch-clamp recording and Channelrhodopsin-2 (ChR2)-assisted circuit mapping, here we show that activation of LC system-derived noradrenergic axon terminals was sufficient to produce an α1- A_R_-EPSC in dorsal raphe serotonin neurons. Further, we show that activation of presynaptic α2-adrenergic receptors (α2-A_R_) inhibited the α1-A_R_-EPSC via inhibition of noradrenaline release. In a subset of serotonin neurons (∼50%), activation of postsynaptic α2-A_R_ produced a small outward current. Collectively, the *in vitro* data suggest that α2-A_R_ inhibit the activity of dorsal raphe serotonin neurons by at least two mechanisms. Indeed, after *in vivo* treatment with an α2-A_R_ agonist, clonidine, fewer serotonin neurons expressed c-fos, an early gene product used as a proxy for neuronal activity [13]. The results reveal that in the dorsal raphe, α2- A_R_ inhibit noradrenaline-dependent synaptic transmission resulting in reduced excitability of the serotonin neurons.

## Materials and Methods

### Animals

Group-housed male and female wild-type C57BL/6J (>2 months old, The Jackson Laboratory, #000664) mice were used for retrograde tracing and electrophysiology experiments except those using optogenetics. For Channelrhodopsin2-assisted circuit mapping, male and female TH-IRES-Cre mice (>2 months old, [14]) were used. All studies were conducted in accordance with the National Institutes of Health Guide for the Care and Use of Laboratory Animals with the approval of the National Institute on Drug Abuse Animal Care and Use Committee or the University of Iowa with the approval of the University of Iowa Institutional Animal Care and Use Committee.

### Retrograde tracing

Mice were anesthetized with ketamine/xylazine (80 mg/kg; 20 mg/kg) and placed in a stereotaxic frame (David Kopf Instruments). Dorsal raphe (AP -4.4; ML 1.19, 20° angle; DV -3.62 mm) microinjections of Fluoro-Gold™ (FG, Fluorochrome LLC, 50 nL, 4%) were made at a rate of 10 nL/min. After one week, mice were euthanized and perfused transcardially with cold PBS followed by 4% paraformaldehyde in PBS (pH 7.4). Brains were removed and stored in 4% paraformaldehyde at 4°C overnight. Coronal sections (50 μm) were made using a vibratome (Leica). Slices was immunostained for TH and anti-fluorogold (1:500, Fluorochrome LLC). For complete methodology, see Supplemental Materials and Methods.

### Optogenetics

Mice were anesthetized with ketamine/xylazine (80 mg/kg; 20 mg/kg) and placed in a stereotaxic frame (David Kopf Instruments). Bilateral microinjections were made into LC (AP -4.9; ML +/- 1; DV -4 mm). 200 nL of a virus encoding for channelrhodopsin-2 (ChR2) and enhanced yellow fluorescent protein (eYFP) (AAV_5_-Ef1α- DIO-ChR2-eYFP, University of North Carolina Gene Therapy Center) was injected at a rate of 100 nL/min. After ∼3 weeks to allow for expression, mice were anesthetized with isoflurane, brains were removed and transferred to modified Krebs’ buffer for brain slice electrophysiology. After recording, brain slices were transferred to 4% paraformaldehyde in PBS for 2 h, then stored in PBS with sodium azide to verify expression of ChR2 by immunostaining TH antibody (1:1000, Aves Labs) to label noradrenaline neurons in the LC. For complete methodology, see Supplemental Materials and Methods.

### Brain slice preparation and electrophysiological recordings

Brain slices and electrophysiological recordings were made as previously described [15]. In brief, mice were deeply anesthetized with isoflurane and euthanized by decapitation. Brains were removed and placed in warmed and bubbled (95/5% O_2_/CO_2_) modified Krebs’ buffer containing (in mM): 126 NaCl, 2.5 KCl, 1.2 MgCl_2_, 1.2 CaCl_2_, 1.2 NaH_2_PO_4_, 21.5 NaHCO_3_, and 11 D-glucose with 5 μM MK-801 to reduce excitotoxicity and increase slice viability. In the same solution, coronal dorsal raphe slices (240 μm) were obtained using a vibrating microtome (Leica) and incubated at 28 °C >30 minutes prior to recording.

Electrophysiological recordings were made from serotonin neurons [12] at 35 °C with Multiclamp 200B and 700B amplifiers (Molecular Devices), Digidata 1440A and 1550B A/D converters (Molecular Devices), and Clampex software (Molecular Devices) with borosilicate glass electrodes (World Precision Instruments) wrapped with Parafilm to reduce pipette capacitance. Pipette resistances were 2.9 to 4.5 MΩ when filled with an internal solution containing, (in mM) 104.56 K-methylsulfate, 3.73 KCl, 5.3 NaCl, 4.06 MgCl_2_, 4.06 CaCl_2_, 7.07 HEPES (K), 3.25 BAPTA (K4), 0.26 GTP (sodium salt), 4.87 ATP (sodium salt), 4.59 creatine phosphate (sodium salt), pH 7.24 with KOH, mOsm ∼274, for whole-cell patch-clamp recordings. Series resistance was monitored throughout the experiment. Reported voltages are corrected for a liquid junction potential of -8 mV between the internal and external solution. All drugs were applied via bath application or iontophoresis. Synaptic currents were evoked on 90-s intervals by applying brief pulses (0.5 ms, 60 Hz) of electrical stimulation to the brain slice via a borosilicate glass monopolar stimulating electrode placed within 200 μm of the recorded neuron in the presence of GluN (MK-801), GluA/GluK (DNQX or NBQX, 3 μM), GABA_A_ (picrotoxin, 100 μM), and 5-HT1A (WAY-100635, 300 nM) receptor blockers to isolate the α1-A_R_-EPSC.

### Protein c-fos immunohistochemistry

Mice received an i.p. injection of clonidine hydrochloride (0.03 mg/kg) or an equal volume of saline and then euthanized 140 min later (Euthasol, i.p.) for transcardial perfusion, brain extraction, post-fixation, and sectioning as described above. Slices were immunostained for c-fos and TPH2. For complete methodology, see Supplemental Materials and Methods.

### Materials

MK-801, NBQX, WAY-100635, prazosin, UK-14.304, noradrenaline, and idazoxan were obtained from Tocris. Clonidine hydrochloride injection was obtained through the University of Iowa Hospital Pharmacy. All other reagents were obtained from Sigma-Aldrich.

### Experimental design and statistical analyses

Data were analyzed using Clampfit 11.1. Data are presented as representative traces, or in scatter plots where each point is an individual cell, and bar graphs with means ± SEM. *n*=number of cells for electrophysiological experiments and *n*=number of mice for immunohistochemical experiments. When possible (within-group comparisons), significant differences were determined for two group comparisons by Wilcoxon matched-pairs signed rank test. Significant mean differences in between-group comparisons were determined by Mann-Whitney tests or two-way ANOVA. A difference of p<0.05 was considered significant. Exact values are reported unless p<0.0001 or >0.999. Statistical analysis was performed using GraphPad Prism 9 (GraphPad Software, Inc.).

## Results

### Noradrenaline afferents in the dorsal raphe arise from the locus coeruleus system

It is well-established that *in vivo* dorsal raphe neurons require noradrenergic input to fire action potentials, but the source of noradrenaline is not founded. Initially, we performed retrograde tracing of Fluoro-Gold™ (FG, [16], n=3 mice) microinjected in the dorsal raphe to the noradrenergic brain nuclei; broadly divided as the locus coeruleus (LC) system (as defined in [17]) and non-LC nuclei in the ventral and dorsomedial medulla. FG was chosen due to more efficient labeling of noradrenaline neurons with FG than other conventional or retrograde viral tracers [18,19]. Noradrenaline neurons were identified by location and immunostaining for tyrosine hydroxylase (TH), the rate-limiting enzyme in the production of dopamine and noradrenaline. On average, FG was found in 94% of TH^+^ neurons in LC proper (555/593 neurons, Fig. 1A-B), 70% of TH^+^ neurons in sub-coeruleus (Sub-C, 334/479 neurons, Fig. 1B) and 70% of TH^+^ neurons in the rostral non-continuous extension of LC (Rnc-LC, 95/137 neurons, Fig. 1B, [17]). In contrast, 18% of TH^+^ neurons in the medulla contained FG (16/90 neurons, Fig. 1B). To test for functional connectivity between the LC system and the dorsal raphe, an optogenetic strategy was employed using the TH-Cre transgenic mouse line [14] that expresses Cre-recombinase in TH^+^ (dopamine and noradrenaline) neurons. TH-Cre mice received bilateral LC microinjections of virus encoding the light-activated cation channel, channelrhodopsin-2 (ChR2) to express ChR2 in noradrenaline neurons, along with enhanced yellow fluorescent protein (eYFP) as a marker of viral transduction (Fig. 1C). ∼3 weeks later, mice were euthanized, and brains were collected for brain slice electrophysiology. Whole-cell current-clamp recordings were made from eYFP^+^ LC neurons. ChR2 was activated with trains of light stimulation (5 or 10 pulses at 60 Hz, 5 ms, 473 nm) which produced action potential firing (Fig. 1D). In the dorsal raphe, whole-cell voltage-clamp recordings were made from serotonin neurons. ChR2-expressing axon terminals were activated with the same trains of light stimulation. In 12 of 20 neurons, the activation of ChR2 was sufficient to produce an α1-A_R_-EPSC that was blocked by an α1-A_R_ inverse agonist, prazosin (Fig. 1E). On average, the amplitude of the optically stimulated α1-A_R_-EPSC was -12.1±2.9 pA (Fig. 1E). The time course of the optically evoked α1-A_R_-EPSC was similar to those produced by electrical stimulation [12] including a time-to-peak of ∼3 s and duration of ∼34 s. Thus, LC noradrenergic axons project to the dorsal raphe and release noradrenaline to produce α1-A_R_-dependent synaptic transmission.

**Figure 1.**
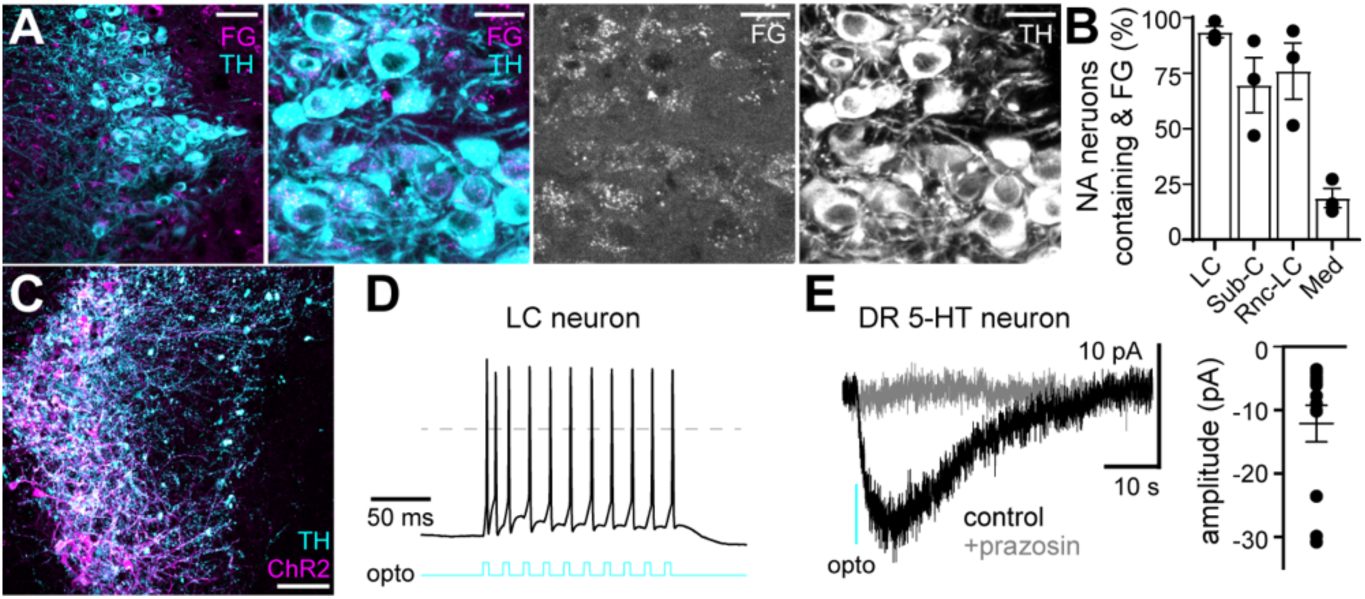
Selective activation of LC noradrenergic axons in the dorsal raphe produces an α1-A_R_-EPSC. A) Representative maximum intensity projection confocal image of noradrenaline neurons in the LC of wild-type mice. Sections were immunostained for tyrosine hydroxylase (TH, cyan) labeling noradrenaline LC neurons and Fluoro-Gold (FG, magenta) demonstrating retrograde transport of FG from the dorsal raphe to LC neurons; scale bars 50 μm (left most) and 20 μm. B) Percentage of TH^+^ neurons that also contained FG in LC, Sub-C, Rnc-LC, and medulla (Med). n=3 mice. C) Representative maximum intensity projection confocal image of noradrenaline neurons in the LC of TH-Cre mice (TH immunostaining, cyan) expressing ChR2-eYFP (ChR2, magenta) after viral transduction following microinjection of AAV_5_-Ef1α-DIO-ChR2-eYFP into the LC; scale bar, 100 μm. D) Representative whole-cell current-clamp recording from an LC neuron expressing ChR2 demonstrating that ChR2 activation (10 pulses at 60 Hz, 5 ms, 473 nm) drove action potential firing. E) Representative whole-cell voltage-clamp recording from a serotonin neuron in the dorsal raphe. Activation of ChR2 expressed in LC noradrenergic axons (train of 5 or 10 pulses at 60 Hz, 5 ms, 473 nm) was sufficient to produce an α1-A_R_-EPSC. Line and error bars represent mean±SEM.

### Activation of α2-adrenergic receptors inhibit noradrenaline-dependent excitatory synaptic transmission

Inhibitory α2-adrenergic receptors (α2-A_R_) are expressed in LC neurons and, when activated, hyperpolarize LC neurons [20–22] and prevent the release of noradrenaline from axon terminals in projection areas [23–30]. To evaluate the consequence of α2-A_R_ activation on noradrenaline-dependent synaptic transmission, whole-cell voltage-clamp recordings were made from dorsal raphe serotonin neurons at 35° C in the presence of GluN, GluA/GluK, GABA_A_, and 5-HT1A receptor antagonists. A train of 5 electrical stimuli (60 Hz) delivered to the brain slice every 90 seconds produced an α1-A_R_-EPSC as previously described [11,12]. Application of an α2-A_R_ agonist, UK-14,304 (0.3 - 3 μM), nearly abolished the α1-A_R_-EPSC (Fig. 2A and 2C, p<0.0001, n=12). Pretreatment of the brain slice with an α2-A_R_ antagonist, idazoxan (1 μM), prevented the reduction by UK-14,304, demonstrating selective action of UK-14,304 on α2-A_R_ (Fig. 2B and 2C, p=0.0003, n=12 and 5). Clonidine (10 µM), a partial α2-A_R_ agonist, also eliminated the α1-A_R_-EPSC (p=0.002, n=6, data not shown). To determine whether activation of α2-A_R_ was inhibiting noradrenaline release, we bypassed the presynaptic element and noradrenaline was focally applied onto the soma via a high-resistance pipette and iontophoresis (300 ms). Iontophoretically applied noradrenaline produced an inward current of ∼ -22 pA (I_NA_, Fig. 2D-F). Application of UK-14,304 (300 nM) had no effect on the amplitude of I_NA_ (Fig. 2D-F, p=0.46, n=8). Thus, activation of α2-A_R_ inhibits the α1-A_R_-EPSC via presynaptic inhibition of noradrenaline release.

**Figure 2.**
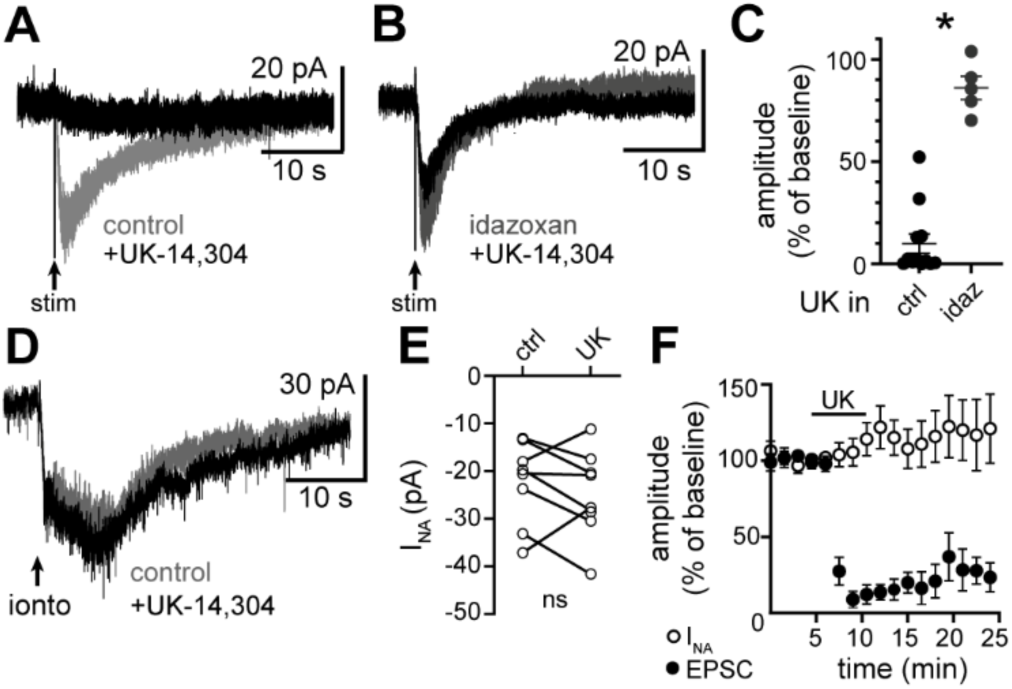
Activation of α2-A_R_ inhibits the α1-A_R_-EPSC via a presynaptic mechanism. A) Representative whole-cell voltage-clamp recording of the α1-A_R_-EPSC in control conditions and in UK-14,304. B) Representative whole-cell voltage-clamp recording of the α1-A_R_-EPSC in an α2-A_R_ antagonist, idazoxan, and in UK-14,304 with idazoxan. C) Plot of the percent remaining of the α1-A_R_-EPSC after application of UK-14,304 in control conditions and in idazoxan (p=0.0003, n=12 and 5, Mann-Whitney test). D) Representative whole-cell voltage-clamp recording of the α1-A_R_-dependent inward current produced by focal iontophoretic application of noradrenaline (ionto) in control conditions and in UK-14,304. E) Plot of the amplitude of the inward current produced by iontophoretic application of NA (I_NA_) in control conditions (ctrl) and in UK-14,304 (UK, p=0.46, n=8, Wilcoxon matched-pairs signed rank test. F) Time course of the inhibition of the α1-A_R_- EPSC (black circles) by UK-14,304, shown in comparison to the lack of effect of UK-14,304 on I_NA_ (white circles). Line and error bars represent mean±SEM. ns denotes not significant, * denotes statistical significance.

### Presynaptic inhibition of noradrenaline release by α2-adrenergic receptors does not depend on membrane potential of innervating axon terminals

To test whether activation of α2-A_R_ inhibited noradrenaline release by activation of voltage-dependent potassium channels, we applied a widely used inhibitor of voltage-gated potassium channels, 4-aminopyridine (4-AP, 100 μM). In the presence of 4-AP, UK-14,304 (1 µM) inhibited the α1-A_R_-EPSC by ∼51%, in contrast to the ∼90% inhibition in control conditions (Fig. 3A-C, p=0.002, n=12 and 7). Thus, blocking voltage-gated potassium channels with 4-AP reduced presynaptic inhibition of noradrenaline release by α2-A_R_. One possible interpretation of these results is that activation of α2-A_R_ increases 4-AP-sensitive potassium conductance, which could hyperpolarize the axon terminal thereby reducing vesicular release of noradrenaline. Alternatively, α2-A_R_ may inhibit voltage-gated calcium channels; an effect that could be partially mitigated by enhanced calcium influx due to 4-AP block of voltage-gated potassium channels and the subsequent broadening of the action potential [31,32]. To distinguish these possibilities, we increased extracellular potassium from 2.5 mM to 10.5 mM which reduces the chemical driving force of potassium and the hyperpolarization from potassium efflux at subthreshold potentials. The reduction in potassium efflux from elevating extracellular potassium was observed as an apparent inward current in the recorded neuron due to a reduction in potassium leak current (Fig. 3D, p=0.016, n=7), which served as a control for solution exchange. Increasing extracellular potassium to 10.5 mM increased the amplitude of the α1-A_R_-EPSC (Fig. 3E, p=0.031, n=6) presumably by increasing noradrenaline release and producing a depolarizing shift in the reversal potential of the α1-A_R_-EPSC [11]. But activation of α2-A_R_ with UK-14,304 in 10.5 mM extracellular potassium still inhibited the α1-A_R_-EPSC robustly, ∼91% (Fig. 3F, p=0.031, n=6). Thus, it is unlikely that presynaptic inhibition of noradrenaline release by α2-adrenergic receptors occurs via increasing potassium conductance.

**Figure 3.**
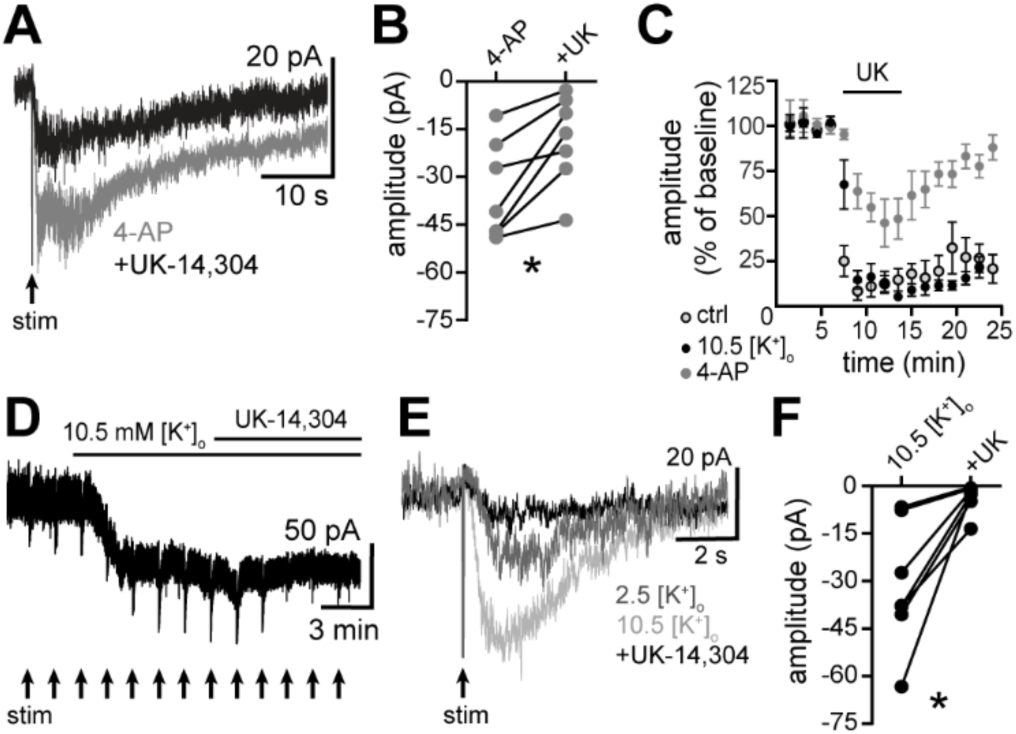
Inhibitory effect of α2-A_R_ activation does not depend on membrane potential of innervating axon terminals. A) Representative whole-cell voltage-clamp recording of the α1-A_R_-EPSC in 4-AP and in UK-14,304 with 4-AP. B) Plot of the amplitude of the α1-A_R_-EPSC in 4-AP and in 4-AP with UK-14,304 (+UK, p=0.016, n=7, Wilcoxon matched-pairs signed rank test). C) Time course of the inhibition of the α1-A_R_-EPSC by UK-14,304 in control conditions (gray circles with black outline), in 10.5 mM extracellular potassium ([K^+^]_o_) (black circles) and in 4-AP (gray circles). Depolarizing innervating axon terminals by increasing [K^+^]_o_ from 2.5 mM to 10.5 mM did not prevent the inhibition of the α1-A_R_-EPSC by activation of α2-A_R_ with UK-14,304; shown in representative traces (D and E) and in grouped data (F, p=0.03, n=6, Wilcoxon matched-pairs signed rank test). Line and error bars represent mean±SEM. * denotes statistical significance.

### Activation of α2-adrenergic receptors hyperpolarizes a subset of serotonin neurons

In previous *in vivo* and rat brain slice studies, a subset of dorsal raphe neurons is hyperpolarized directly by noradrenaline [5,33] via activation of α2-A_R_ opening membrane ion channels [34]. In agreement, we found that UK-14,304 produced an outward current (∼18 pA) in 52% of serotonin neurons (“responders”, Fig. 4A-B) which was associated with a decrease in membrane resistance (p=0.029, n=16, Fig. 4C), indicative of opening of ion channels. In the remaining 48%, there was no change in current to UK-14,304 detected over baseline noise (“non-responders”, Fig. 4A-B) and no change in membrane resistance (p=0.45, n=15, Fig. 4C). When applied in 10.5 mM extracellular potassium which shifts the reversal potential for potassium close to the holding potential (V_hold_: -65 mV, calculated E_K_: -66 mV), UK-14,304 failed to produce an outward current (2.6±5.2 pA, p=0.56, n=6, data not shown). Taken together these data indicate that a subset of serotonin neurons express α2-A_R_ that when activated, produce an outward current likely through activation of potassium channels.

**Figure 4.**
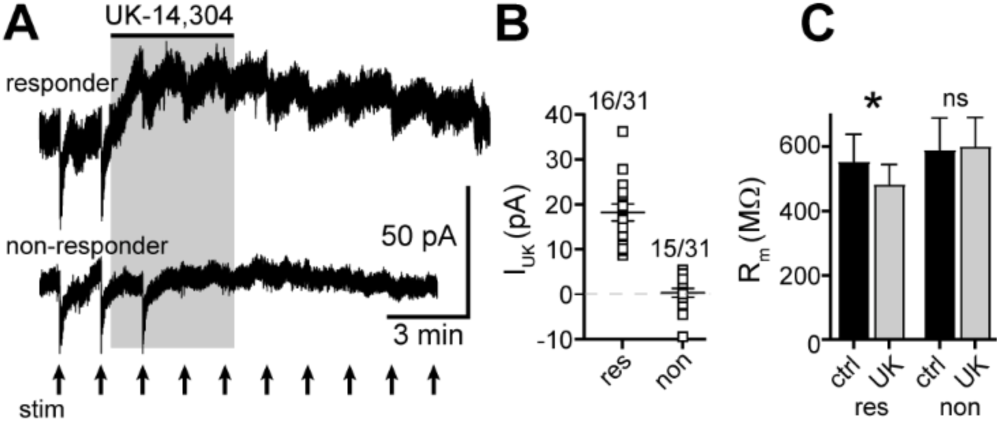
Activation of α2-A_R_ produces an outward current in a subset of serotonin neurons. A) Representative whole-cell voltage-clamp recordings of the outward current produced by UK-14,304 in a subset of neurons (responders) while other neurons were unaffected (non-responders). We note that the inhibition of the α1-A_R_-EPSC (evoked at each arrow) to UK-14,304 was observed in both responders and non-responders. B) Plot of the amplitude of the UK-14,304-induced outward current in responders (res) and non-responders (non). C) UK-14,304 produced a small decrease in membrane resistance, indicative of opening of ion channels, in responders (res, p=0.029, n=16, Wilcoxon matched-pairs signed rank test) but not non-responders (non, p=0.45, n=15, Wilcoxon matched-pairs signed rank test). Line and error bars represent mean±SEM. ns denotes not significant, * denotes statistical significance.

### *In vivo* activation of α2-adrenergic receptors reduces c-fos expression in dorsal raphe serotonin neurons

Collectively, our *in vitro* data suggest that activation of α2-A_R_ in the dorsal raphe has two forms of inhibitory action on serotonin neurons: 1) by eliminating α1-A_R_-dependent excitation and 2) by directly hyperpolarizing a subset of neurons. To evaluate the action of α2-A_R_ activation *in vivo*, we utilized the expression of the immediate early gene c-fos as a proxy for neuronal activity [13,35]. Male and female mice received a single i.p. injection of clonidine (0.03 mg/kg) or an equal volume of saline and perfused 140 minutes later. Their brains were collected for immunostaining of dorsal raphe serotonin neurons for c-fos and tryptophan hydroxylase 2 (TPH2), as marker of serotonin neurons [12]. Counts were made of midline serotonin neurons across the rostral-caudal and ventral-dorsal axes of the dorsal raphe (Fig. 5A). We found that fewer serotonin neurons expressed c-fos after clonidine treatment when compared to saline-treated mice (Fig. 5B, p=0.032, n=18 mice in each condition). When disaggregated by sex, there was still a significant effect of clonidine treatment that was comparable between males and females (Fig. 5C, main effect of treatment, p=0.022; main effect of sex, p=0.37; interaction, p=0.93; n=9 mice for each group). Together, these results are consistent with the *in vitro* data and indicate that *in vivo* activation of α2-A_R_ reduces the activity of dorsal raphe serotonin neurons.

**Figure 5.**
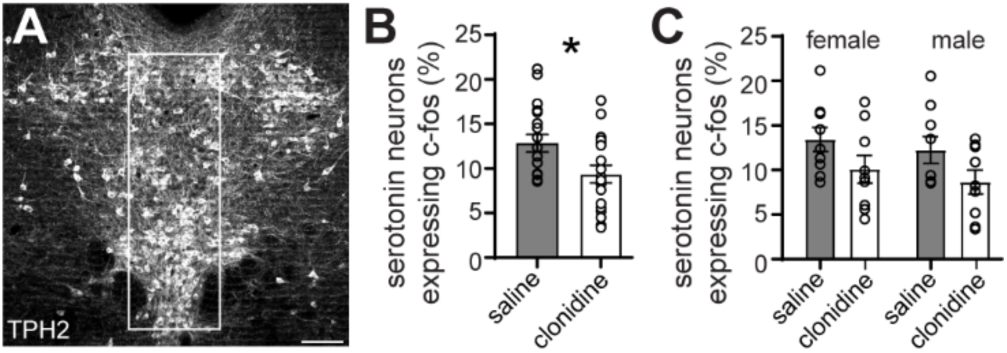
*In vivo* activation of α2-A_R_ reduces c-fos expression in dorsal raphe serotonin neurons. A), Representative maximum intensity projection confocal image of the dorsal raphe. Sections were immunostained for tryptophan hydroxylase (TPH2, grey) labeling the serotonin neurons, scale bar: 100 μm. The box indicates the midline area where analyses were performed from 40ξ images. B and C) A single i.p. injection of clonidine reduced the expression of c-fos in serotonin neurons (p=0.032, n=18 mice/group); in both male and female mice (Two-way ANOVA: main effect of treatment, p=0.022; main effect of sex, p=0.37; interaction, p=0.93; n=9 mice/group). Line and error bars represent mean±SEM. * denotes statistical significance.

## Discussion

### Resolving a source of noradrenaline to the dorsal raphe

*In vivo*, dorsal raphe serotonin neurons are driven to fire action potentials via excitatory noradrenergic input. Depletion of noradrenaline or microinjection of α1-A_R_ antagonists in the dorsal raphe silences the neurons [3–5,8] and decreases serotonin release [9]. In contrast, *in vivo* microinjection of α1-A_R_ agonists increases action potential firing in the majority of dorsal raphe neurons [4,33] and elevates serotonin release [6] demonstrating that α1-A_R_ signaling is a fundamental bidirectional modulator of the excitability and release of serotonin from dorsal raphe serotonin neurons. Electron microscopic work shows that the serotonin neurons are directly innervated by noradrenergic terminals [36], but the source of noradrenaline afferents has been highly debated. Early research that utilized mechanical or chemical lesioning of LC demonstrated a subsequent loss of noradrenaline-containing terminals in the dorsal raphe ([37,38], but also see [39]). In addition, administration of the neurotoxin DSP-4, which leads to the destruction of LC system-derived axon terminals and spares non-LC noradrenergic axons (reviewed in [40,41]), dramatically reduces the level of noradrenaline in dorsal raphe homogenates [42] providing compelling evidence that LC projects to the dorsal raphe. However, this work was challenged by *in vivo* recordings which demonstrated that electrical stimulation of the LC fails to change the firing rate of dorsal raphe neurons [43,44]. Further, retrograde labeling from the dorsal raphe to noradrenergic neurons with horseradish peroxidase or cholera-toxin subunit B typically produced only sparse labeling in LC and the other noradrenergic or adrenergic brain regions [44–48], suggesting that the dorsal raphe may be innervated weakly by many noradrenergic brain regions. Alternatively, these conventional tracers may not be readily endocytosed by the noradrenergic terminals (e.g. [18]). Here using FG, we show significant retrograde labeling from the dorsal raphe to all of the LC system, including LC proper, sub-coeruleus, and Rnc-LC. A smaller sub-population of noradrenaline neurons in the medulla were also labeled. Selective optogenetic activation of LC system-derived noradrenergic axons caused the release of noradrenaline and produced an α1-A_R_-EPSC in dorsal raphe serotonin neurons. Collectively, these data indicate that the LC system is the largest contributor to noradrenaline release in the dorsal raphe without excluding the possibility of some noradrenergic afferents arising from the medulla.

### Dual inhibitory action of α2-adrenergic receptors in the dorsal raphe

Using microdialysis, electrochemical detection, and radiolabeling, it is known that α2-adrenergic receptors inhibit noradrenaline release from peripheral sympathetic nerves [26,49] and from LC cell bodies [23,50,51] and axon terminals in the brain [23,25–29]. But the degree to which α2-A_R_-mediated inhibition of release reduces noradrenaline-dependent synaptic transmission is not well-described. In the present study, we found that activation of α2-A_R_ with exogenous agonists, UK-14,304 or clonidine, eliminated the α1-A_R_-EPSC in dorsal raphe serotonin neurons via presynaptic inhibition of noradrenaline release. The data are consistent with prior *in vivo* studies. Activation of α2-A_R_ with clonidine, whether administered systemically or into the LC, silences action potential firing in LC neurons and consequently, most of the dorsal raphe serotonin neurons [8]. Conversely, systemic or intra-raphe administration of an α2-A_R_ antagonist accelerates action potential firing in the serotonin neurons [52–54]. The ability of α2-A_R_ activation to inhibit noradrenaline release did not depend on the membrane potential of the innervating axon terminals nor was likely due to activation of potassium channels, despite the observation that blocking voltage-gated potassium channels with 4-AP reduced presynaptic inhibition of noradrenaline release by α2-A_R_. While the mechanism remains to be fully elucidated, the prediction is that activated α2-A_R_ reduce calcium influx into the terminals, as observed in LC cell bodies [55], thereby reducing vesicular release through inhibition of voltage-gated calcium channels. Then 4-AP can functionally counteract the inhibition of voltage-gated calcium channels by broadening the action potential and extending the open-time of the remaining calcium channels. Alternatively, activated α2-A_R_ may inhibit noradrenaline release in a manner that is independent of calcium influx [56].

Here we also report evidence of postsynaptic α2-A_R_ on a subset of serotonin neurons. In contrast to the intense labeling of α2A mRNA in LC, *in situ* hybridization shows a small amount of mRNA for the α2C subtype in the dorsal raphe [57], although there is pharmacological evidence for functional α2A-A_R_ [58]. Activation of the postsynaptic α2-A_R_ produced a small outward current in ∼50% of the neurons likely due to activation of potassium conductance. Our results are consistent with previous *in vivo* and rat brain slice studies that report a minority (∼13-40%) of dorsal raphe neurons are hyperpolarized by noradrenaline ([3,33], but see [5]) via activation of α2-A_R_ opening ion channels [8,34]. Our estimates may be higher due to the selective recording from serotonin neurons, as opposed to surrounding GABA and dopamine neurons [12,59], or since we utilized bath application of an α2-A_R_ agonist which would activate all receptors irrespective of their location relative to the soma. Indeed, *in vivo* when a high level of noradrenaline is applied to dorsal raphe neurons, the inhibitory action of noradrenaline dominates in up to 88% of the neurons [5], which may suggest that α2-A_R_ are located on distal dendrites. We note that despite evidence of a postsynaptic α2-A_R_-dependent outward current to exogenous agonist, an α2-A_R_-mediated inhibitory postsynaptic current (α2-A_R_-IPSC) evoked by synaptic release of noradrenaline was not observed. In some instances throughout the brain, receptors are only located at extrasynaptic sites and ‘spill-over’ of neurotransmitter outside of the synaptic cleft is required for their activation and the generation of the postsynaptic response [60,61]. However, in the dorsal raphe, the spatial spread of noradrenaline is controlled tightly by transporter-dependent reuptake [12]. There is little evidence of spill-over of noradrenaline unless reuptake is impaired [12]. Thus, it may be that postsynaptic α2-A_R_ on serotonin neurons are located at only extrasynaptic sites and require ‘spill-over’ of noradrenaline to generate an IPSC.

### Summary and health implications

Dysregulation of the noradrenaline and serotonin systems have been linked to pathological forms of hyper- and hypo-arousal that present in ADHD, obsessive-compulsive, bipolar, and schizophrenia, as well as idiopathic and stress-induced sleep disturbances, pain syndromes, PTSD, depression, and anxiety disorders. The current lack of understanding of how the serotonin and noradrenaline systems interact at the level of neural circuits and synapses limits the development of better therapies.

We have previously shown in brain slices that synaptic activation of α1-A_R_ depolarizes serotonin neurons and drives action potential firing [11]. The temporal and spatial spread of extracellular noradrenaline is limited by efficient reuptake via noradrenaline transporters [12]. Here we show that LC noradrenergic axons project to the dorsal raphe and release noradrenaline to produce α1-A_R_-dependent synaptic transmission. α2- adrenergic ‘autoreceptors’ on presynaptic axon terminals, likely the α2A subtype [51,57,62], reduce the excitability of dorsal raphe serotonin neurons by eliminating α1-A_R_-dependent excitation. In addition, the excitability of a subset of serotonin neurons is expected to be reduced further with activation of α2-adrenergic ‘heteroreceptors’, likely the α2A or α2C subtype [57,58], on the membrane through activation of potassium conductance. Lastly, *in vivo* activation of α2-A_R_ by systemic administration of clonidine reduced the expression of c-fos in the dorsal raphe serotonin neurons, indicating reduced neural activity. Thus, α2-A_R_ are critical regulators of serotonin neuron excitability, which expected to reduce serotonin release. There are prior studies that administration of an α2-A_R_ agonist into the dorsal raphe reduces serotonin release through action on pre- and postsynaptic α2-A_R_ [6,9,58] which may be through reduced excitability, direct inhibition of voltage-gated calcium channels [63], and reductions in the biosynthesis of serotonin [64].

Pharmacotherapies that target the noradrenaline system, either by preventing transporter-dependent reuptake (antidepressants or psychostimulants) or directly targeting α-adrenergic receptors (agonists or antagonists, reviewed in [65]) are used commonly to treat neuropsychiatric conditions with features of hypo- and hyper-arousal. Numerous studies using human post-mortem brain have implicated the over-expression or super-sensitivity of α2-A_R_ in the etiology of major depressive disorder [66–68], reviewed in [69] Currently, the first-line treatment for major depressive disorder are serotonin-selective reuptake inhibitors (SSRIs), but upwards of 30% of patients do not experience symptom relief from SSRIs [70]. Based on the results of the present study, over-expression of α2-A_R_ may lead to profound reductions in serotonin release, which ultimately may contribute to SSRI-resistant depression. This hypothesis is supported by prior work that demonstrates that the over-expression and altered function of α2-A_R_ in platelets taken from untreated patients with major depressive disorder is predictive of treatment outcome [71].

## Acknowledgements

A portion was supported by the Intramural Research Program at the National Institute on Drug Abuse (H.S.H and S.C.G). The opinions expressed in this article are the authors’ own and do not reflect the views of the NIH/DHHS. The present address for H.S.H. is in the Department of Psychology, University of Washington, Seattle, Washington, 98195, USA. We would like to thank Drs. Yeka Aponte (NIH) and Brenton Laing (University of Mississippi) for providing materials and expertise for the retrograde labeling experiments. We would also like Nora O’Prey, Addison Eckard, Jeff Rudman, Kathryn Cochrane, and Andrew Kain for their technical assistance in c-fos immunohistochemistry and analysis.

## Author contributions

Aleigha Gugel: Substantial contributions to the acquisition, analysis, or interpretation of data for the work; Drafting the work or revising it critically for important intellectual content; Final approval of the version to be published; and Agreement to be accountable for all aspects of the work in ensuring that questions related to the accuracy or integrity of any part of the work are appropriately investigated and resolved.
Erik A. Ingebretsen: Substantial contributions to the acquisition, analysis, or interpretation of data for the work; Final approval of the version to be published; and Agreement to be accountable for all aspects of the work in ensuring that questions related to the accuracy or integrity of any part of the work are appropriately investigated and resolved.
Holly S. Hake: Substantial contributions to the acquisition, analysis, or interpretation of data for the work; Final approval of the version to be published; and Agreement to be accountable for all aspects of the work in ensuring that questions related to the accuracy or integrity of any part of the work are appropriately investigated and resolved.
Stephanie C. Gantz: Substantial contributions to the conception or design of the work; or the acquisition, analysis, or interpretation of data for the work; Drafting the work or revising it critically for important intellectual content; Final approval of the version to be published; and Agreement to be accountable for all aspects of the work in ensuring that questions related to the accuracy or integrity of any part of the work are appropriately investigated and resolved.

## SUPPLEMENTARY MATERIALS AND METHODS

### Retrograde tracing

Slices were blocked and permeabilized in PBS with 0.2% Triton X-100 and 5% normal donkey serum (Jackson ImmunoResearch) for 2 h. Immunohistochemistry was then performed by using chicken anti-TH antibody (1:1000, Aves Labs) overnight. After washing, sections were further incubated in an Alexa Fluor 594 or 647 donkey anti-chicken (1:500) in PBS for 2 h. FG-injected tissue was incubated in anti-fluorogold (1:500, Fluorochrome LLC) in PBS overnight. Sections were then washed, mounted, and coverslipped with Fluoromount-G™, a DAPI mounting medium (ThermoFisher Scientific). Fluorescent images (20ξ and 40ξ) were obtained with FV1000 confocal microscope system (Olympus). Neurons that were both TH^+^ and either FG^+^ or FG^-^ were counted using FIJI software.

### Optogenetics

Slices were blocked and permeabilized in PBS containing 10% normal donkey serum (Jackson ImmunoResearch) and 0.5% Triton X-100 for 5 h then incubated overnight in chicken anti-TH antibody (1:1000, Aves Labs) to label noradrenaline neurons in the LC. After washing, slices were incubated in Alexa Fluor 594 donkey anti-chicken (1:500) for 2 h. Sections were then washed, mounted, and coverslipped with Fluoromount-G™. Fluorescent images were obtained with FV1000 confocal microscope system (Olympus).

### c-fos immunohistochemistry

Slices were washed in PBS, permeabilized and blocked in PBS with 0.5% Triton-X and 10% normal donkey serum for 3 h. Slices were incubated overnight in rabbit anti-c-Fos (Cell Signaling, 1:4000), washed in PBS, and incubated in donkey anti-rabbit Alexa Fluor 647 (Jackson Immunoresearch, 1:1000) for 2 h. Then slices were washed (PBS) and incubated in goat anti-TPH2 (Abcam, 1:1000) overnight, washed, and then incubated in donkey anti-goat Alexa Fluor 488 (Jackson Immunoresearch, 1:1000) for 2 h. Slices were washed, mounted, and coverslipped with Fluoromount-G™. Fluorescent images (40ξ magnification) were obtained with an Axiovert 100 confocal microscope system (Zeiss). Expression of c-fos and TPH2 immunostaining was quantified by manual counting using FIJI software by investigators that were blinded to the sex and treatment group. Counts were made of midline TPH2^+^ neurons across the rostral-caudal and ventral-dorsal axes of the dorsal raphe.

## REFERENCES

1. Aston-Jones G, Rajkowski J, Cohen J. Role of locus coeruleus in attention and behavioral flexibility. Biol Psychiatry. 1999;46:1309–1320.

2. Ågren H, Koulu M, Saavedra JM, Potter WZ, Linnoila M. Circadian covariation of norepinephrine and serotonin in the locus coeruleus and dorsal raphe nucleus in the rat. Brain Res. 1986;397:353–358.

3. Baraban JM, Wang RY, Aghajanian GK. Reserpine suppression of dorsal raphe neuronal firing: mediation by adrenergic system. Eur J Pharmacol. 1978;52:27–36.

4. Baraban JM, Aghajanian GK. Suppression of serotonergic neuronal firing by α-adrenoceptor antagonists: Evidence against GABA mediation. Eur J Pharmacol. 1980;66:287–294.

5. Baraban JM, Aghajanian GK. Suppression of firing activity of 5-HT neurons in the dorsal raphe by alpha-adrenoceptor antagonists. Neuropharmacology. 1980;19:355–363.

6. Pudovkina OL, Cremers TIFH, Westerink BHC. Regulation of the release of serotonin in the dorsal raphe nucleus by α_1_ and α_2_ adrenoceptors. Synapse. 2003;50:77–82.

7. Pudovkina OL, Cremers TIFH, Westerink BHC. The interaction between the locus coeruleus and dorsal raphe nucleus studied with dual-probe microdialysis. Eur J Pharmacol. 2002;445:37–42.

8. Svensson TH, Bunney BS, Aghajanian GK. Inhibition of both noradrenergic and serotonergic neurons in brain by the α-adrenergic agonist clonidine. Brain Res. 1975;92:291–306.

9. Bortolozzi A, Artigas F. Control of 5-Hydroxytryptamine Release in the Dorsal Raphe Nucleus by the Noradrenergic System in Rat Brain. Role of α-Adrenoceptors. Neuropsychopharmacology. 2002;28:421– 434.

10. Vandermaelen CP, Aghajanian GK. Electrophysiological and pharmacological characterization of serotonergic dorsal raphe neurons recorded extracellularly and intracellularly in rat brain slices. Brain Res. 1983;289:109–119.

11. Gantz SC, Moussawi K, Hake HS. Delta glutamate receptor conductance drives excitation of mouse dorsal raphe neurons. eLife. 2020;9.

12. Khamma JK, Copeland DS, Hake HS, Gantz SC. Spatiotemporal Control of Noradrenaline-Dependent Synaptic Transmission in Mouse Dorsal Raphe Serotonin Neurons. J Neurosci. 2022;42:968–979.

13. Sheng M, Greenberg ME. The regulation and function of c-*fos* and other immediate early genes in the nervous system. Neuron. 1990;4:477–485.

14. Lindeberg J, Usoskin D, Bengtsson H, Gustafsson A, Kylberg A, Söderström S, et al. Transgenic expression of Cre recombinase from the tyrosine hydroxylase locus. Genesis. 2004;40:67–73.

15. Copeland DS, Gugel A, Gantz SC. Potentiation of neuronal activity by tonic GluD1 current in brain slices. EMBO Rep. 2023;24.

16. Saleeba C, Dempsey B, Le S, Goodchild A, McMullan S. A Student’s Guide to Neural Circuit Tracing. Front Neurosci. 2019;13.

17. Grzanna R, Molliver ME. The locus coeruleus in the rat: an immunohistochemical delineation. NSC. 1980;5:21–40.

18. Delfs JM, Zhu Y, Druhan JP, Aston-Jones GS. Origin of noradrenergic afferents to the shell subregion of the nucleus accumbens: anterograde and retrograde tract-tracing studies in the rat. Brain Res. 1998;806:127– 140.

19. Ganley RP, Werder K, Wildner H, Zeilhofer HU. Spinally projecting noradrenergic neurons of the locus coeruleus display resistance to AAV2retro-mediated transduction. Mol Pain. 2021;17:1–10.

20. Williams JT, Henderson G, North RA. Characterization of α_2_-adrenoceptors which increase potassium conductance in rat locus coeruleus neurones. NSC. 1985;14:95–101.

21. Williams JT, Marshall KC. Membrane Properties and Adrenergic Responses in Locus Coeruleus Neurons of Young Rats. J Neurosci. 1987;7:3687–3694.

22. Courtney NA, Ford CP. The Timing of Dopamine- and Noradrenaline-Mediated Transmission Reflects Underlying Differences in the Extent of Spillover and Pooling. J Neurosci. 2014;34:7645–7656.

23. Mateo Y, Pineda J, Meana JJ. Somatodendritic α_2_-Adrenoceptors in the Locus Coeruleus Are Involved in the In Vivo Modulation of Cortical Noradrenaline Release by the Antidepressant Desipramine. J Neurochem. 1998;71:790–798.

24. Washburn M, Moises HC. Electrophysiological Correlates of Presynaptic α_2_-Receptor-Mediated Inhibition of Norepinephrine Release at Locus Coeruleus Synapses in Dentate Gyrus. J Neurosci. 1989;9:2131–2140.

25. Wang B, Wang Y, Wu Q, Huang H, Li S. Effects of α2A Adrenoceptors on Norepinephrine Secretion from the Locus Coeruleus during Chronic Stress-Induced Depression. Front Neurosci. 2017;11:1–8.

26. Trendelenburg AU, Klebroff W, Hein L, Starke K. A study of presynaptic α_2_-autoreceptors in α_2A/D_-, α_2B_- and α_2C_-adrenoceptor-deficient mice. Naunyn-Schmiedeberg’s Arch Pharmacol. 2001;364:117–130.

27. Veldhuizen MJA van, Feenstra MGP, Heinsbroek RPW, Boer GJ. *In vivo* microdialysis of noradrenaline overflow: effects of α-adrenoceptor agonists and antagonists measured by cumulative concentration-response curves. Br J Pharmacol. 1993;109:655–660.

28. L’Heureux R, Dennis T, Curet O, Scatton B. Measurement of Endogenous Noradrenaline Release in the Rat Cerebral Cortex In Vivo by Transcortical Dialysis: Effects of Drugs Affecting Noradrenergic Transmission. J Neurochem. 1986;46:1794–1801.

29. Bücheler MM, Hadamek K, Hein L. Two α_2_-adrenergic receptor subtypes, α_2A_ and α_2C_, inhibit transmitter release in the brain of gene-targeted mice. NSC. 2002;109:819–826.

30. Frankhuijzen AL, Wardeh G, Hogenboom F, Mulder AH. Alpha 2-adrenoceptor mediated inhibition of the release of radiolabelled 5-hydroxytryptamine and noradrenaline from slices of the dorsal region of the rat brain. Naunyn-Schmiedeberg’s Arch Pharmacol. 1988;337:255–260.

31. Sabater VG, Rigby M, Burrone J. Voltage-Gated Potassium Channels Ensure Action Potential Shape Fidelity in Distal Axons. J Neurosci. 2021;41:5372–5385.

32. Bean BP. The action potential in mammalian central neurons. Nat Rev Neurosci. 2007;8:451–465.

33. Couch JR. Responses of neurons in the raphe nuclei to serotonin, norepinephrine and acetylcholine and their correlation with an excitatory synaptic input. Brain Res. 1970;19:137–150.

34. Yoshimura M, Higashi H, Nishi S. Noradrenaline mediates slow excitatory synaptic potentials in rat dorsal raphe neurons in vitro. Neurosci Lett. 1985;61:305–310.

35. Gantz SC, Ortiz MM, Belilos AJ, Moussawi K. Excitation of medium spiny neurons by “inhibitory” ultrapotent chemogenetics via shifts in chloride reversal potential. eLife. 2021;10.

36. Baraban JM, Aghajanian GK. Noradrenergic innervation of serotonergic neurons in the dorsal raphe: Demonstration by electron microscopic autoradiography. Brain Res. 1981;204:1–11.

37. Loizou LA. Projections of the nucleus locus coeruleus in the albino rat. Brain Res. 1969;15:563–566.

38. Chu N-S, Bloom FE. The catecholamine-containing neurons in the cat dorsolateral pontine tegmentum: Distribution of the cell bodies and some axonal projections. Brain Res. 1974;66:1–21.

39. Roizen MF, Jacobowitz DM. Studies on the origin of innervation of the noradrenergic area bordering on the nucleus raphe dorsalis. Brain Res. 1976;101:561–568.

40. Bortel A. Nature of DSP-4 induced neurotoxicity. In: Kostrzewa R, editor. Handbook of Neurotoxicity. Springer, New York, NY. 2014.

41. Ross SB, Stenfors C. DSP4, a Selective Neurotoxin for the Locus Coeruleus Noradrenergic System. A Review of Its Mode of Action. Neurotox Res. 2015;27:15–30.

42. Cassano T, Gaetani S, Morgese MG, Macheda T, Laconca L, Dipasquale P, et al. Monoaminergic Changes in Locus Coeruleus and Dorsal Raphe Nucleus Following Noradrenaline Depletion. Neurochem Res. 2009;34:1417–1426.

43. Wang RY, Gallager DW, Aghajanian GK. Stimulation of pontine reticular formation suppresses firing of serotonergic neurones in the dorsal raphe. Nature. 1976;264:365–368.

44. Anderson CD, Pasquier DA, Forbes WB, Morgane PJ. Locus coeruleus-to-dorsal raphe input examined by electrophysiological and morphological methods. Brain Res Bull. 1977;2:209–221.

45. Aghajanian GK, Wang RY. Habenular and other midbrain raphe afferents demonstrated by a modified retrograde tracing technique. Brain Res. 1977;122:229–242.

46. Peyron C, Luppi PH, Fort P, Rampon C, Jouvet M. Lower Brainstem Catecholamine Afferents to the Rat Dorsal Raphe Nucleus. J Comp Neurol. 1996;364:402–413.

47. Kim M-A, Lee HS, Lee BY, Waterhouse BD. Reciprocal connections between subdivisions of the dorsal raphe and the nuclear core of the locus coeruleus in the rat. Brain Res. 2004;1026:56–67.

48. Luppi P-H, Aston-Jones G, Akaoka H, Chouvet G, Jouvet M. Afferent projections to the rat locus coeruleus demonstrated by retrograde and anterograde tracing with cholera-toxin B subunit and *Phaseolus vulgaris* leucoagglutinin. NSC. 1995;65:119–160.

49. Altman JD, Trendelenburg AU, MacMillan L, Bernstein D, Limbird L, Starke K, et al. Abnormal Regulation of the Sympathetic Nervous System in α_2A_-Adrenergic Receptor Knockout Mice. Mol Pharmacol. 1999;56:154–161.

50. Palij P, Stamford JA. Real-time monitoring of endogenous noradrenaline release in rat brain slices using fast cyclic voltammetry: 1. Characterisation of evoked noradrenaline efflux and uptake from nerve terminals in the bed nucleus of stria terminalis, pars ventralis. Brain Res. 1992;587:137–146.

51. Callado LF, Stamford JA. α_2A_-But not α_2B/C_-adrenoceptors modulate noradrenaline release in rat locus coeruleus: voltammetric data. Eur J Pharmacol. 1999;366:35–39.

52. Garratt JC, Crespi F, Mason R, Marsden CA. Effects of idazoxan on dorsal raphe 5-hydroxytryptamine neuronal function. Eur J Pharmacol. 1991;193:87–93.

53. Freedman JE, Aghajanian GK. Idazoxan (RX 781094) selectively antagonizes α_2_-adrenoceptors on rat central neurons. Eur J Pharmacol. 1984;105:265–272.

54. Haddjeri N, Blier P, Montigny C de. Effect of the *Alpha*-2 Adrenoceptor Antagonist Mirtazapine on the 5-Hydroxytryptamine System in the Rat Brain. J Pharmacol Exp Ther. 1996;277:861–871.

55. Ingram S, Wilding TJ, McCleskey EW, Williams JT. Efficacy and Kinetics of Opioid Action on Acutely Dissociated Neurons. Mol Pharmacol. 1997;52:136–143.

56. Schwartz DD. Activation of *Alpha*-2 Adrenergic Receptors Inhibits Norepinephrine Release by a Pertussis Toxin-Insensitive Pathway Independent of Changes in Cytosolic Calcium in Cultured Rat Sympathetic Neurons. J Pharmacol Exp Ther. 1997;282:248–255.

57. McCune SK, Voigt MM, Hill JM. Expression of multiple alpha adrenergic receptor subtype messenger RNAs in the adult rat brain. NSC. 1993;57:143–151.

58. Hopwood SE, Stamford JA. Noradrenergic modulation of serotonin release in rat dorsal and median raphé nuclei via α_1_ and α_2A_ adrenoceptors. Neuropharmacology. 2001;41:433–442.

59. Huang KW, Ochandarena NE, Philson AC, Hyun M, Birnbaum JE, Cicconet M, et al. Molecular and anatomical organization of the dorsal raphe nucleus. eLife. 2019;8.

60. Otis TS, Mody I. Differential activation of GABAA and GABAB receptors by spontaneously released transmitter. J Neurophysiol. 1992;67:227–235.

61. Beenhakker MP, Huguenard JR. Astrocytes as Gatekeepers of GABA_B_ Receptor Function. J Neurosci. 2010;30:15262–15276.

62. Lee A, Rosin DL, Van Bockstaele EJ. α_2A_-adrenergic receptors in the rat nucleus locus coeruleus: subcellular localization in catecholaminergic dendrites, astrocytes, and presynaptic axon terminals. Brain Res. 1998;795:157–169.

63. Li Y-W, Bayliss DA. Activation of α_2_-adrenoceptors causes inhibition of calcium channels but does not modulate inwardly-rectifying K^+^ channels in caudal raphe neurons. NSC. 1997;82:753–765.

64. Yoshioka M, Matsumoto M, Togashi H, Smith CB, Saito H. α_2_-Adrenoceptor modulation of 5-HT biosynthesis in the rat brain. Neurosci Lett. 1992;139:53–56.

65. Langer SZ. α_2_-Adrenoceptors in the treatment of major neuropsychiatric disorders. Trends Pharmacol Sci. 2015;36:196–202.

66. Ordway GA, Widdowson PS, Smith KS, Halaris A. Agonist Binding to α_2_-Adrenoceptors Is Elevated in the Locus Coeruleus from Victims of Suicide. J Neurochem. 1994;63:617–624.

67. Ordway GA, Schenk J, Stockmeier CA, May W, Klimek V. Elevated agonist binding to α_2_-adrenoceptors in the locus coeruleus in major depression. Biol Psychiatry. 2003;53:315–323.

68. González AM, Pascual J, Meana JJ, Barturen F, Arco CD, Pazos A, et al. Autoradiographic Demonstration of Increased α_2_-Adrenoceptor Agonist Binding Sites in the Hippocampus and Frontal Cortex of Depressed Suicide Victims. J Neurochem. 1994;63:256–265.

69. Cottingham C, Wang Q. α_2_ adrenergic receptor dysregulation in depressive disorders: Implications for the neurobiology of depression and antidepressant therapy. Neurosci Biobehav Rev. 2012;36:2214–2225.

70. McIntyre RS, Alsuwaidan M, Baune BT, Berk M, Demyttenaere K, Goldberg JF, et al. Treatment-resistant depression: definition, prevalence, detection, management, and investigational interventions. World Psychiatry. 2023;22:394–412.

71. Gurguis GNM, Vo SP, Griffith JM, Rush AJ. Platelet alpha_2A_-adrenoceptor function in major depression: G_i_ coupling, effects of imipramine and relationship to treatment outcome. Psychiatry Res. 1999;89:73–95.

